# Microbial populations are shaped by dispersal and recombination in a low biomass subseafloor habitat

**DOI:** 10.1101/2021.02.03.429647

**Authors:** Rika E. Anderson, Elaina D. Graham, Julie A. Huber, Benjamin J. Tully

## Abstract

The subseafloor is a vast habitat that supports microorganisms that have a global scale impact on geochemical cycles. Many of the endemic microbial communities inhabiting the subseafloor consist of small populations under growth-limited conditions. For small populations, stochastic evolutionary events can have large impacts on intraspecific population dynamics and allele frequencies. These conditions are fundamentally different from those experienced by most microorganisms in surface environments, and it is unknown how small population sizes and growth-limiting conditions influence evolution and population structure in the subsurface. Using a two-year, high-resolution environmental time-series, we examine the dynamics of microbial populations from cold, oxic crustal fluids collected from the subseafloor site North Pond, located near the mid-Atlantic ridge. Our results reveal rapid shifts in overall abundance, allele frequency, and strain abundance across the time points observed, with evidence for homologous recombination between coexisting lineages. We show that the subseafloor aquifer is a dynamic habitat that hosts microbial metapopulations that disperse frequently through the crustal fluids, enabling gene flow and recombination between microbial populations. The dynamism and stochasticity of microbial population dynamics in North Pond suggests that these forces are important drivers in the evolution of microbial populations in the vast subseafloor habitat.

**Significance Statement:** The cold, oxic subseafloor is an understudied habitat that is difficult to access, yet important to global biogeochemical cycles and starkly different compared to microbial habitats on the surface of the Earth. Our understanding of microbial evolution and population dynamics is largely molded by studies of microbes living in surface habitats that can host 10-1,000 times more microbial biomass than is frequently observed in the subsurface. This study provides an opportunity to observe population dynamics within a low biomass, growth-limited environment and reveals that microbial populations in the subseafloor are influenced by changes in selection pressure and gene sweeps. In addition, recombination between strains that have dispersed from elsewhere within the aquifer has an important impact on the evolution of microbial populations. Much of the microbial life on the planet exists under growth-limited conditions and the subseafloor provides a natural laboratory to explore how life evolves in such environments.

---

**T**he marine subsurface is home to more than half of the archaea and bacteria inhabiting the oceans (1, 2). Marine subsurface habitats are frequently energy-limited, which impacts the total biomass that can be supported in these environments (3). As a result, microbial communities in these habitats are often composed of relatively low biomass (4–7) (*<* 10^4^ cells per unit volume). Population size can have important consequences for how microbial populations evolve over time, and factors such as genetic drift can have an exaggerated impact on small microbial populations compared to habitats in which microbial populations are larger. Most previous studies of adaptation in natural microbial communities have focused on habitats with high biomass and thus larger population sizes, such as the surface ocean, deep-sea hydrothermal vents and freshwater lakes (8–11). It is unclear how natural selection and genetic drift shape the evolution of microbial communities in low-biomass habitats, despite the fact that a substantial fraction of global microbial biomass is found in such habitats.

North Pond, located near the mid-Atlantic ridge (12), offers a unique opportunity to examine dynamics of microbial adaptation in natural populations in a low-biomass, growth-limited habitat. The North Pond site consists of exposed crustal ridges with a depression that has accumulated *<* 100 m of low permeability sediment, causing the bulk of fluid exchange and transport to occur beneath the sediment basin (13). This sub-seafloor aquifer connects the deep waters of the oceans through a porous network of crustal rock, which globally contains about 2% of the oceans’ volume (14–16). The crustal aquifer beneath the sediments of North Pond is accessible through two circulation obviation retrofit kits (CORKs) installed by the IODP ocean drilling program in 2011, providing direct access to the fluids circulating within the aquifer, while preventing exchange with the overlaying water and sediment environments (17, 18). Repeated sampling in 2012, 2014, and 2017, including an eight sample in situ time-series collected from 2012-2014, revealed a dynamic microbial community with a high degree of functional redundancy in a complex hydrological system depleted in dissolved organic carbon (DOC) (6, 19–21). The aquifer microbial communities are distinct from bottom seawater, and there are multiple lines of genetic evidence to suggest that the microbial community within the aquifer is motile and metabolically flexible, with the ability to carry out both autotrophic and organotrophic pathways (6, 19, 22). The North Pond subseafloor microbial community has low biomass (10^3^-10^4^ cells ml^-1^) and incubation experiments suggest the North Pond microbial community is growth-limited and that increasing temperature and enrichment with carbon sources results in rapid growth (6, 7, 22).

A major question in subsurface habitats is what the major drivers of evolution are in energy- and growth-limited environments such as North Pond. It has been hypothesized that in energy- and growth-limited environments, such as low-deposition deep-sea sediments, adaptive mutations would be unable to sweep through populations and mutations would accumulate as a result of energy limitation (23). However, many crustal habitats like North Pond do not reflect the limited fluid flow and isolated cellular environments in low-energy sediments, and instead may reflect the population dynamics of soils with high fluid connectivity (24). North Pond fluids are capable of traversing distances of approximately (21) 2-40 m day^-1^, though this fluid flux is spatially and temporally complex, resulting in convective and oscillatory fluid movement rather than linear flow along the North-South axis (13). Fluid movement appears to impact the microbial community, as previous analysis of the system revealed discrete ecological units within the time-series that would oscillate between presence and absence with extended time periods (*e*.*g*., 100s of days) between reoccurrences (6). However, it is unclear how this dynamic environment impacts the endemic microbial populations.

Intraspecies diversity results from the introduction of variation through mutation or gene flow, and that variation is removed through selection and genetic drift. Thus, examination of patterns of genomic variation within populations can provide insights into evolutionary dynamics within a system. Here, we present an analysis of the intraspecific population dynamics of the North Pond aquifer microbial community from a metagenomic time series spanning 10 time points over a two-year period. We sought to characterize the dynamics of population variation over time in order to determine what processes constrain variation within microbial populations at North Pond, and to assess the balance between deterministic processes such as selection and more stochastic processes such as dispersal. We show that abundant microorganisms within the crustal fluids have patterns of genome diversity suggesting that large-scale genomic sweeps are infrequent and small gene sweeps, recombination, and dispersal of populations from the larger metapopulation within the aquifer may have a more pronounced impact on genome diversity and therefore evolution in this habitat. These results provide new understanding of microbial population dynamics over time and how microorganisms adapt and persist in the largest habitat on the planet.

## Results and Discussion

### Improving North Pond metagenome-assembled genomes

By comparing and combining the total set of new MAGs to the original 2018 NORP MAGs (6) (*n* = 195 MAGs), it was possible to replace 22 2018 NORP MAGs with newly reconstructed MAGs with higher completion and/or lower redundancy statistics and to produce 137 completely new MAGs, previously unrecovered from the metagenomic datasets (Supplemental Data 1). Additional results and discussion of these MAG results can be found in the Supplementary Information.

### Targeting of specific populations in North Pond fluids

To conduct a robust analysis of genome dynamics at borehole U1382A over the two-year time series, we focused on MAGs that occurred in at least three samples with ≥ 5 RPKM (Table 1; Figure 1; Supplemental Data 2). In total, six MAGs met these criteria. In addition, we identified another four MAGs with high coverage (*>* 30 RPKM) in only one sample. In all four instances, these MAGs had low coverage (*<* 3 RPKM) in the other time points, with three MAGs having extremely low coverage at other time points (*<* 0.15 RPKM) (Table 1; Figure 1; Supplemental Data 2). The set of mapped metagenomic reads to each of the 10 MAGs represents the overall metapopulation diversity of a mixture of strains that represent a microbial population at a specific site at a specific point in time. Therefore, the terms “MAG” and “population” are used interchangeably or in conjunction with each other for the remainder of the manuscript. We conducted an in-depth analysis of these 10 MAGs to understand the impacts of dispersal, selection, and recombination on microbial populations in the aquifer. Due to the complexity of fluid movement within the aquifer, it is challenging to determine whether rapid changes in cell abundance result from a dispersal event, whereby cells are flushed from the sampling location, or the result of “rapid” death and growth over the measured time points. Given the dynamism of the aquifer, we assumed that rapid shifts in population structure were likely associated with dispersal; however, given the small population sizes within these microbial communities, drift during bottleneck events remains likely.

**Table 1.**
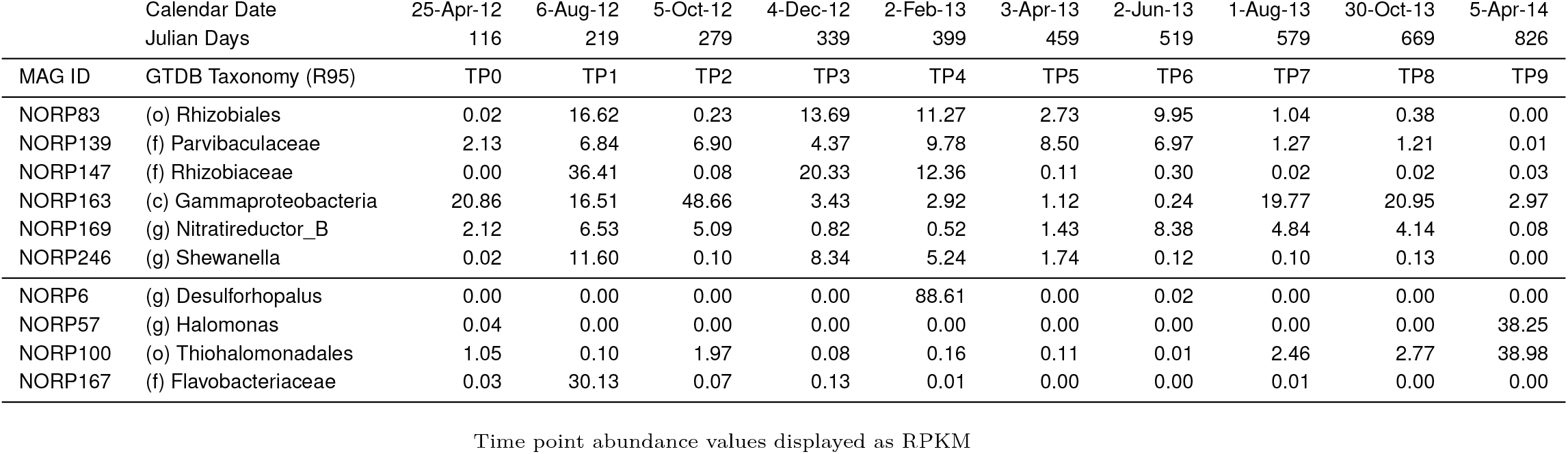
Information for the MAGs of interest.

**Fig. 1.**
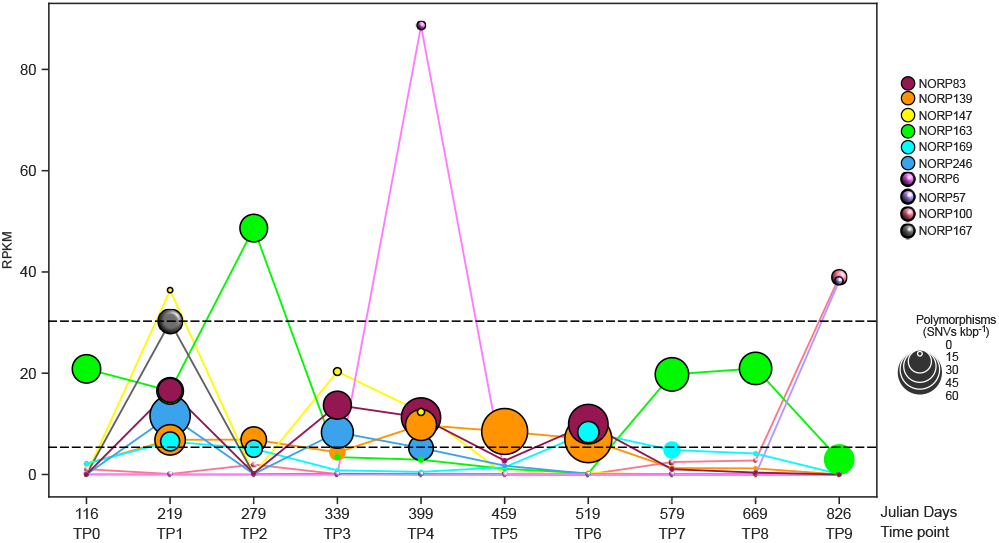
Scatter plot of the abundance for each of the ten MAGs of interest sampled from North Pond CORK U1382A. The size of each marker is scaled to the number of polymorphisms detected in the sample. Dotted lines highlighted RPKM values of 30 and 5, which were used as cutoffs to determine time points of interests. Markers with ≥ 5 RPKM are outlined with black lines.

### MAGs represent extant populations in the aquifer

In general, the planktonic component of the microbial community within the aquifer has low cell density (10^3^ − 10^4^ cell mL^-1^) (6, 7, 22). Under these low biomass conditions, we hypothesized that some of the microbial populations represented by the MAGs may be derived from organisms that had undergone a recent clonal expansion. A clonal expansion occurs when a population undergoes rapid growth, usually due to expansion into a new niche, and subsequently does not have enough time to accumulate mutations within the population (25). All MAGs were assessed for features indicative of a recent clonal expansion. Each of the MAGs had rapid shifts in abundance, and we were particularly interested in the four MAGs (NORP6, NORP57, NORP100, and NORP167) that demonstrated a sudden increase in abundance at a single time point, with extremely low abundance at other time points (Table 1; Figure 1). The number of detected polymorphisms (SNVs kbp^-1^), nucleotide diversity, and evidence of active replication were used as metrics to determine if any of the populations were candidates for clonal expansion (Supplemental Data 3; Supplemental Data 4). Based on genome coverage trough-to-peak (T:P) ratios, most MAGs in the time points of interest had evidence of active replication, with between 1.3-6.8 average genome copies per cell. Despite drastic changes in abundance between time points (increasing from ∼0 to 38.3 and 88.6 RPKM respectively (Table 1)), NORP6 and NORP57 did not return a T:P ratio (Supplemental Data 3), indicating a lack of observable replication. Clonal expansions have been rarely observed in nature. A previous example, from a freshwater environment, observed a population with 0.003 SNVs kbp^-1^ polymorphisms (26). Three of the North Pond MAGs (NORP6, NORP57, and NORP147) had notably few polymorphisms, ranging from 0.2-1.3 SNVs kbp^-1^, but all others had more (5.7-63.8 SNVs kbp^-1^; Figure 1; Supplemental Data 3). Based on the observed nucleotide diversity of the MAGs across all relevant time points (*π* = 0.002 − 0.01), each population is diverse and shares minor alleles (Supplemental Data 3). This level of population diversity is approximately equivalent to that observed for soils in grassland meadows and higher than single species-dominated photosynthetic mats in a connected watershed (11, 24). The accumulation of diversity within the populations of interest, as measured through nucleotide diversity, suggests that there were no large-scale genome sweeps shortly before sampling. Collectively, this evidence suggests that none of the populations that met the criteria for analysis were candidates of recent clonal expansion. Instead, we observed high underlying diversity indicative of a complex system with coexisting subpopulations, approximately equivalent to subspecies or strains.

### Nucleotide-level heterogeneity reveals coexistence of multiple strains within populations

For the six MAGs with high abundance across multiple time points, it was possible to compare patterns in allele frequency and reconstruct strain abundance patterns, revealing the coexistence of multiple subpopulations/strains. The MAG populations represented by NORP169 and NORP246 are estimated to contain approximately three strains each (Figure 2; Figure S1). There were pronounced shifts in the relative abundance of the three strains between time points. For example, in the NORP246 population, strain Str1 is ∼65% of the population in TP1, while in TP3 strain Str2 becomes the dominant strain at ∼51% of the population and reaches ∼62% in TP4 (Figure 2A). The shifts in relative strain abundance were reflected in the major allele frequency, where time points dominated by a single strain have more sites that are fixed at a single allele (TP1 and TP3 major allele frequency 83.1% and 83.4%, respectively), while time points with multiple strains in similar proportions tend to have major allele frequencies closer to 0.5 (TP4 and TP6 major allele frequency 62.5% and 64.6%, respectively; Figure 2C).

**Fig. 2.**
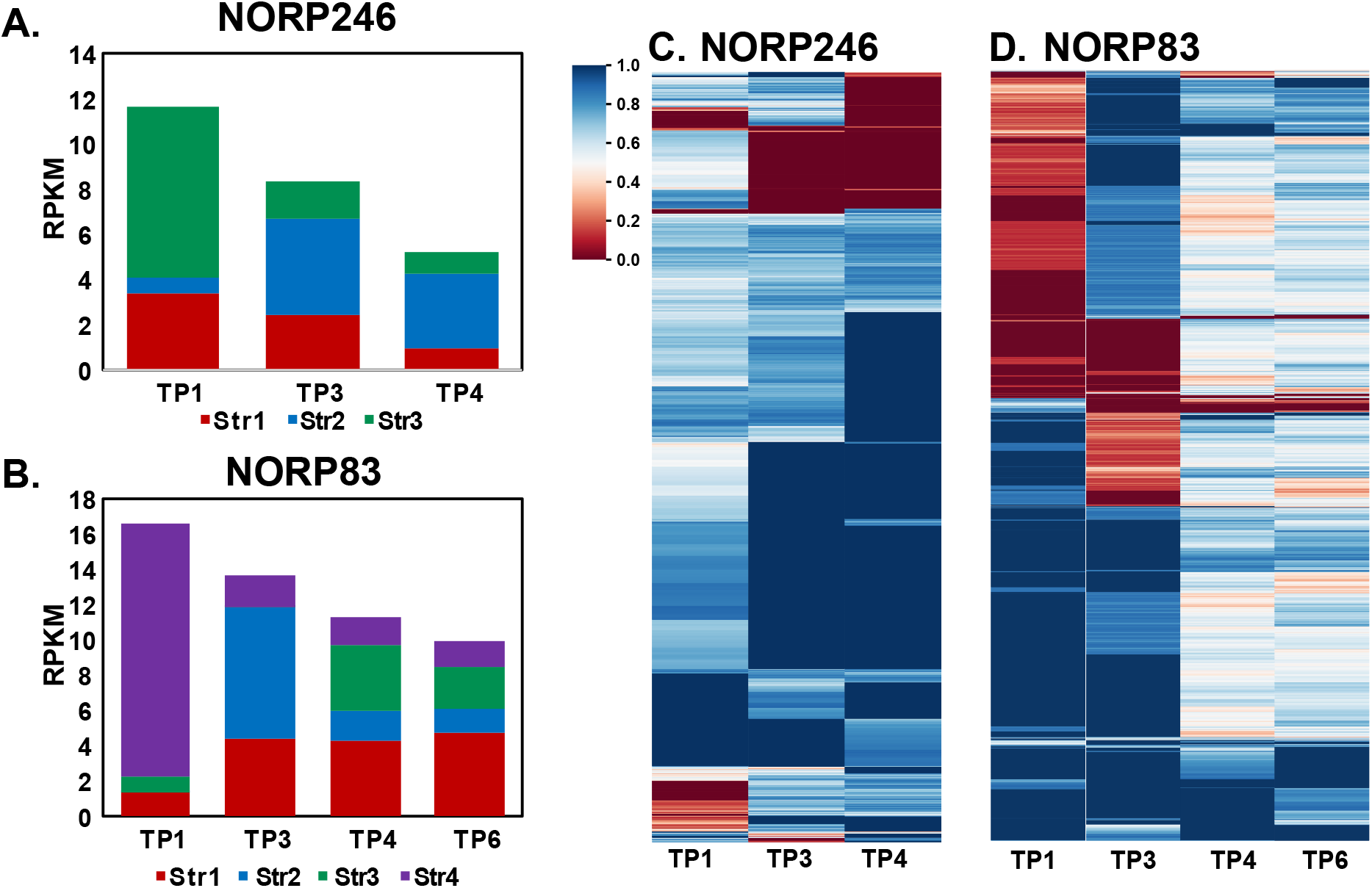
Strain relative abundance as determined using DESMAN for (A) NORP246 and (B) NORP83. Major allele frequency for SNVs detected in time points of interest for (C) NORP246 and (B) NORP83. The major allele frequencies have been hierarchically clustered and scaled from 0-1. Both 0.0 and 1.0 represent a fixed allele; 0.5 represents a split allele.

The other four MAG populations (NORP83, NORP139, NORP147, and NORP163) differed in that during the time points of interest, at least one strain enters the population that was not present at other time points. For example, the NORP139 population consisted of six strains (Str1-Str6; Figure S1). The NORP139 strain Str1 was dominant in TP2 (∼62% of the population), but essentially absent (*<* 0.7%) for other time points. Distinctly, NORP147, in the first time point of interest, was dominated by a single strain, with two different strains appearing as a minor component (*<* 9%) of the population over subsequent time points (Figure S1). For all four of these populations, the introduction of these strains supports the existence of a larger metapopulation for which specific members were observed at North Pond at discrete time points. We observed shifts in the major allele frequency corresponding to the appearance of these new strains. This was exemplified by the transition in strain membership for NORP83 from TP1 to TP3, where the dominant strain Str4 (∼87% of the population) was supplanted by the introduction of strain Str2 (∼54% of the population), and more than half of the alleles shift from being nearly fixed at one allele to another different allele state (Figure 2B & 2D). Moreover, when all four strains are present in TP4 and TP5 in a plurality (14-48% of the population), most of the major allele frequencies are close to 0.5 (Figure 2C & 2D). The ecology of these populations, with multiple strains disappearing and appearing across time points and changes in major allele frequencies corresponding to these shifts, and the high underlying genetic diversity, without indications of recent population-wide removal of diversity, supports an argument that the larger aquifer metapopulation is the source of genetic diversity observed at the North Pond site. This high observed diversity suggests that these strains originate through evolutionary selection and drift elsewhere in the aquifer over time periods longer than those observed in this study.

### Homologous recombination is common and varies by population

If population differentiation occurs elsewhere and throughout the aquifer, the degree of recombination between coexisting strains has important implications for speciation and gene transfer. Low rates of recombination could lead to sympatric speciation or more frequent genome-wide sweeps, whereas high rates of recombination between strains could hold lineages together and also lead to gene-specific sweeps and facilitate gene transfer. We used the decay rate of linkage disequilibrium and the four-gamete test as a proxy for detecting recombination within the populations represented by each MAG. All of the populations had four alleles (denoted as H4, *i*.*e*., the presence of the AB, ab, Ab, and aB haplotypes) at some of the linked biallelic SNVs sites (mean H4 frequencies for each population 0.44%-14.84%). Several of the MAG populations (NORP83, NORP147, and NORP169) had H4 frequencies (0.44%, 1.17%, and 1.45%, respectively) below the lowest values estimated for single species photosynthetic mats (3.4% H4 frequency (11); Table 2). NORP147 did not display decay of the linkage disequilibrium values as genetic distance increased for any of the mutation type pairs, indicating a lack of recombination within the population (S-S, N-S, or N-N; Figure 3A). NORP83 displayed linkage disequilibrium decay (Figure 3B), though the rate of decay was substantially lower than in the other North Pond populations (*e*.*g*., NORP246; Figure 3C; Figure S2), which is an indication of lower recombination rates. NORP57 did not display linkage decay (Figure 3D), but had a slightly larger H4 frequency (5.40%). Both NORP83 and NORP169 had sufficient data to produce an estimate of recombination-to-mutation (gamma/mu; *γ/µ*) from the tool mcorr, which had values (*NORP* 83_*γ/µ*_: 1.10-4.59 and *NORP* 169_*γ/µ*_: 4.71-8.25) similar to that of human pathogens (24, 27) (Supplemental Data 5). Collectively, this evidence suggests the NORP83, NORP57, NORP147, and NORP169 populations did not undergo high rates of recombination relative to other North Pond populations. For NORP147, the limited number of coexisting strains would reduce the opportunity for recombination, but for NORP83 and NORP169, which have multiple coexisting strains, this may imply that intrinsic biological characteristics prevented high rates of recombination within these populations. Diminished rates of recombination could increase the potential for sympatric speciation and genome-wide sweeps.

**Table 2.**
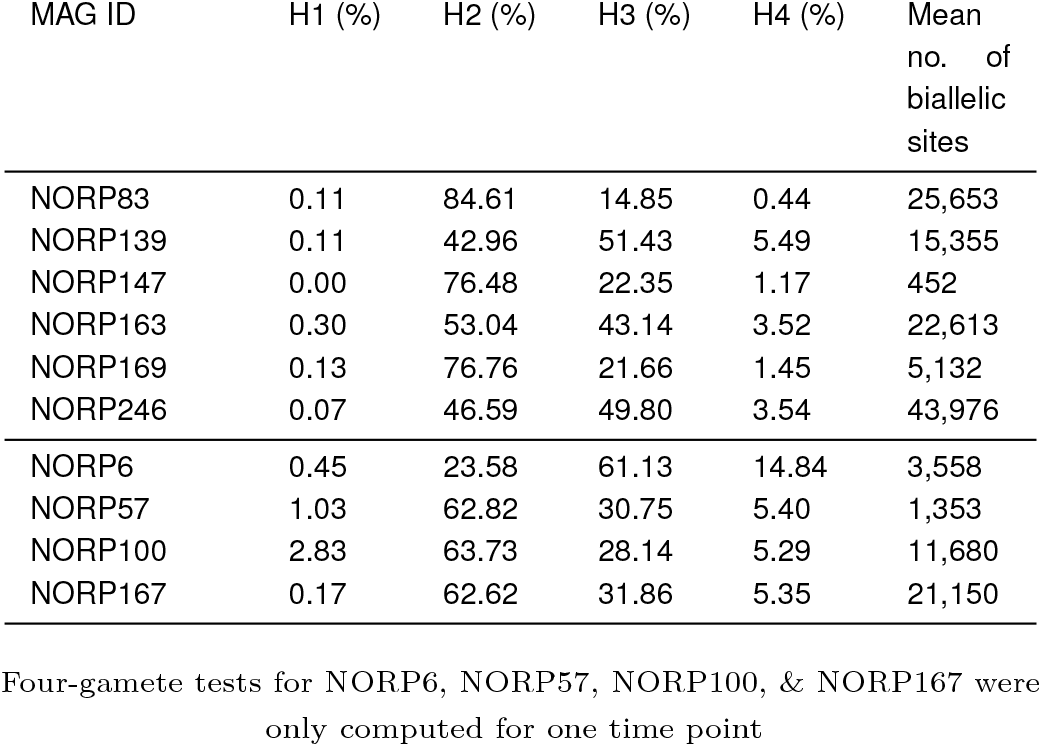
Percent mean four-gamete test results for all time points of interest.

**Fig. 3.**
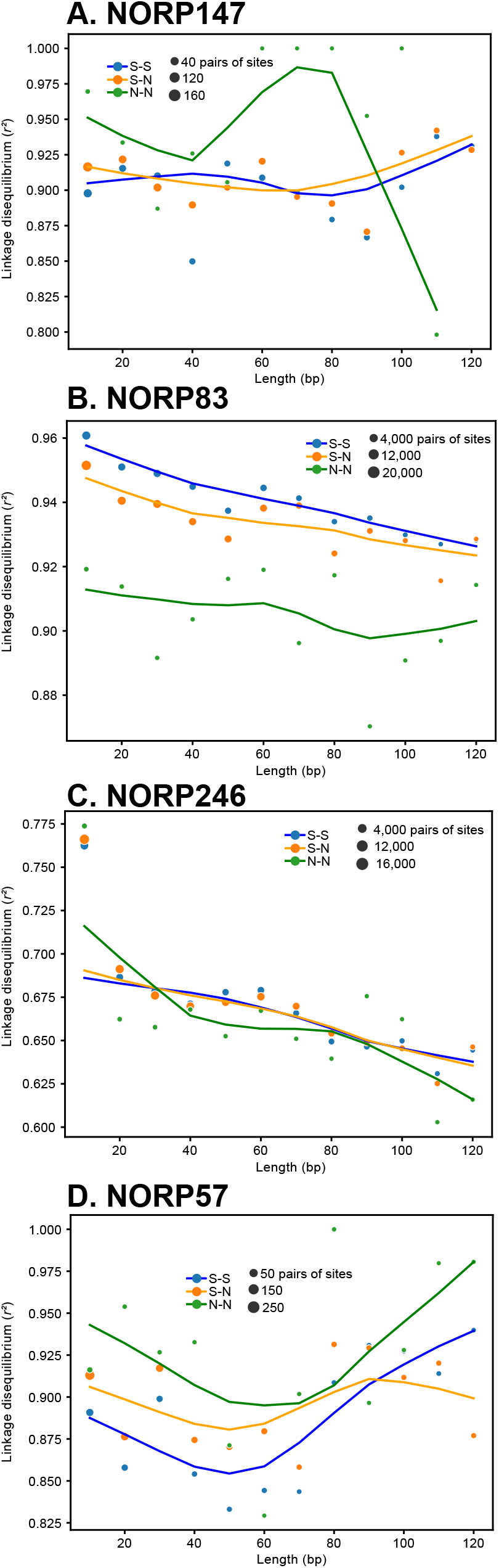
Linkage disequilibrium of *r*^*2*^ for linked SNVs pairs for (A) NORP147, (B) NORP83, (C) NORP246, and (D) NORP57. Each circle is the mean *r*^*2*^ for pairs of linked SNVs at that distance range (*e*.*g*., 1-10 bp, 11-20 bp, *etc*.), with the area proportional to the number of linked SNVs in the mean. Linked SNVs are denoted by their predicted mutation type (nonsynonymous: N, synonymous: S).

Conversely, the other six MAG populations had distinct patterns of linkage decay and higher frequencies of the H4 allele, suggesting likely recombination; however, the degree of linkage disequilibrium varied between them (Table 2; Figure S2). For all of these populations, homologous recombination could maintain cohesion among the coexisting strains, preventing sympatric speciation while potentially facilitating gene-specific sweeps and possibly gene exchange among closely related strains. The balance between recombination and selection dictates the degree to which coexisting microbial lineages can speciate in the absence of geographical barriers (28–30). Recombination can play a cohesive role by maintaining sequence clusters and preventing speciation, particularly if the effects of recombination are lower than that of mutation (31). If recombination rates are low, clusters of organisms can form from selective sweeps, resulting in “ecotypes” that are adapted to specific niches (32). These ecotypes are maintained by periodic selection for or against specific mutations. However, if recombination rates are high enough, specific genes could sweep through populations without initiating a genome-wide sweep (26, 28). Our results suggest that recombination is common but populations likely do not reach panmictic equilibrium in the aquifer, in which recombination occurs frequently enough such that alleles are not linked across microbial genomes. Thus, both genome-wide and gene-specific sweeps are possible among these microbial populations.

### Gene content variations across strains

Genes that are unique to individual strains may provide a selective advantage to each population. Genes limited to specific strains were identified by calculating changes in gene copy number based on observed changes in coverage over time (≥1× change in coverage between time points; Supplemental Data 6). The NORP147 population, dominated by a single strain at all three time points of interest, did not have genes with variable copy number; and NORP139, which had higher comparative rates of recombination, only had five genes with variable gene copy number, for which four had no KEGG annotations. We examined the four other populations to understand the putative functions of the genes that had variable copy numbers over time.

For the NORP169 population, none of the seven variable genes had KEGG annotations, but several of the functional assignments from GenBank implicated transcriptional regulation possibly related to plasmid replication (Figure 4A). Four of the genes were co-localized within the MAG (NORP169 Gene IDs 1205-1208) and shifted from *<* 1× copies per genome in TP1 and TP6 to ∼4× copies per genome in TP2. This small shift corresponded with an increase in strain Str1 and may reflect an extrachromosomal element.

**Fig. 4.**
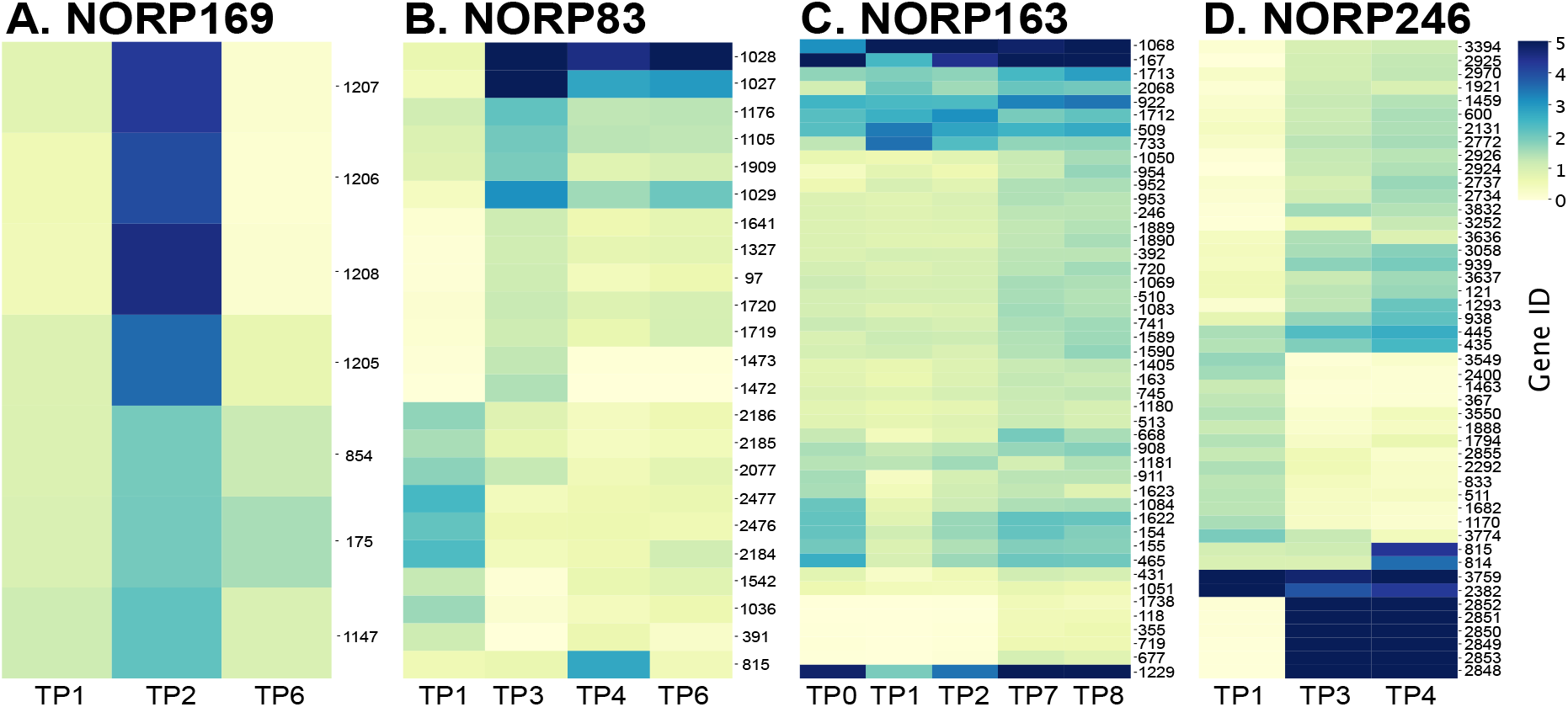
Hierarchically clustered heat map of gene frequencies for genes ≥ 1× change in coverage over the time points of interest for (A) NORP169, (B) NORP83, (C) NORP163, and (D) NORP246. Scaled range from 0 − 5× copies. Maximum value of gene frequency in (D) NORP246 is 21.7× copies. Data available in Supplemental Data 6.

In NORP83, 13 of the 23 variable genes (57%) had no KEGG annotation (Supplemental Data 6). We observed shifts in gene abundance likely associated with the gradual loss of strain Str4 and the introduction of strain Str2 (Figure 2C & 4B). Two sets of genes (NORP83 Gene IDs 2184-2186 and 2476-2477) decreased from ∼2× copies per genome in TP1 to *<* 1× copies in TP3 and were annotated as part of a type IV secretion system (IDs 2184-2186) and chromosome partitioning (ID 2477), which may indicate a role in horizontal gene transfer (33). Conversely, two sets of genes were absent in TP1 (IDs 1472-1473, a putative sodium:proton antiporter and serine hydrolase; IDs 1719-1720, sugar permease) and had *>* 1× copies per genome in TP3 (Figure 4B; Supplemental Data 5). A set of genes (IDs 1027-1029) present at *<* 0.5× copies per genome in TP1 and annotated as putative transposases increased in gene copy in TP3 (5-9× copies per genome) before dropping to an elevated level in TP4 and TP6 (2-6× copies per genome; Figure 4B). These patterns of gene frequency suggest that these transposases were introduced into the North Pond NORP83 population with the arrival of strain Str2.

The NORP163 population had 45 putative genes that changed in gene copy number over time (Figure 4C; Supplemental Data 6). While 29 of the variable genes (64%) had no KEGG annotations, 12 of these genes were putative transposases, with several exceeding *>* 4× copies per genome in all of the time points (Figure 4C). There were five genes that were absent in TP0-TP2 and had *>* 0× copies per genome in TP7-TP8 that were annotated as ribosomal proteins (IDs 167 and 2068), hypothetical proteins (IDs 922 and 1712), and a sulfite reductase (ID 1714; Supplemental Data 6). A larger set of genes changed from *<* 1× copies per genome to *>* 1× copies from TP0 to TP8 (Figure 4C). Co-localized genes of this larger set had functions related to core carbon metabolism (Gene IDs 1083-1084), heavy metal sensing and transcriptional response (IDs 1589-90), and osmolyte transport (IDs 1622-1623). These genes that increased in copy number represent putative functions associated with environmental sensing and response through regulation and substrate transport.

Shifts in gene abundance for the population represented by NORP246 were highly dynamic over time and accompanied by a change in strain abundance, most notably the increase of strain Str2 over time (Figure 2A & 4D). Forty-seven putative genes changed substantially in frequency over three time points in the NORP246 population (Figure 4D). These genes can be divided into three broad groups: (1) present in TP1 and absent in TP3 and TP4, (2) absent in TP1 and present in TP3 and TP4, and (3) absent in TP1 and present in high coverage in TP3 and TP4. These most likely reflect genes that were either present or absent in strain Str2. The 13 putative genes in group one were present in TP1 (∼1× copies per genome) and absent in TP3 and TP4, likely representing genes that were absent in strain Str2. These were predominantly annotated as hypothetical proteins with some conserved domains (*n* = 7; Figure 4D). Three of the putative genes had roles in transport: xanthine permease (ID 1682), sodium/glutamate symporter (ID 833), and cobalt ATP binding cassette-type permease (ID 2292; Supplemental Data 6). Two co-localized genes in this group were annotated as a transposase and reverse transcriptase (IDs 3549-3550; Supplemental Data 6). Conversely, the 23 putative genes in group two were absent in TP1 (*<* 1× copies per genome) and present in TP3 and TP4 (1-2× copies per genome), likely reflecting genes that were present in strain Str2 and absent in strains Str1 and Str3. These were predominantly annotated as hypothetical proteins with conserved domains (*n* = 18; Figure 4D). Genes in group three, which were absent in TP1 and present in high coverage in TP3 and TP4 (15-20× copies per genome), likely representing genes in high abundance in strain Str2, were co-localized (IDs 2848-2853) and lack annotations, except for gene ID 2851 with homology to proteins in GenBank annotated as DNA replication protein (Figure 4D; Supplemental Data 6). This segment appeared to contain a fragment of DNA that in TP3 and TP4 could be independently replicating relative to the NORP246 genome, possibly a plasmid or similar extrachromosomal element. The lack of annotations makes it impossible to determine the exact roles of these genes within the TP3- and TP4-populations, but it is possible that selection favored the functional attributes of strain Str2 over that of strains Str1 and Str3.

There were three general trends in the types of genes that had gene copy variation: (1) genes lacking KEGG annotations, for which function remains unknown; (2) genes annotated as and associated with extrachromosomal replication (*e*.*g*., plasmids, integrases, etc.) and horizontal gene transfer; and (3) genes annotated as transporters. While variation in gene copy numbers over time for genes with functional annotations were identified, we cannot state with certainty that these genes provide a selective advantage for a particular variant, but some functions could provide specialization at the intraspecific level. For example, transporters play an important role in substrate availability and acquisition. The correlation between changes in gene frequency and the introduction/removal of a strain from the North Pond populations reinforces the role of dispersal as the likely mechanism for introducing genomic variation between strains of the same population. These populations may compete for and/or maintain partially overlapping ecological niches, such that changes in gene content may reflect the generation of “ecotypes” amongst the co-occurring strains. In general, the introduction of new strains did not correspond to an increase in total population abundance, perhaps reinforcing the idea that the intraspecific populations shared an ecological niche rather than directly competing for it (34). NORP83, NORP163, and NORP246 had multiple transposases that shifted in copy number over time. An increase in transposase activity has previously been identified as a signature of genetic drift (35). However in this instance, we observed an increase in the number of reads mapping to genes identified as transposes, though it is unclear whether this represented the spreading of transposases to different strains or an increase in transposase copy number within a strain. Other genes that changed in abundance, such as putative plasmids and the type IV secretion system, represent “selfish gene” elements that can propagate between strains during mixing and recombination events. Potentially in a system like North Pond, where growth is slow and infrequent, the actions and dispersal of self propagating elements may play an important role in gene transfer events as they occur outside canonical windows of genomic replication.

### Signatures of gene-specific sweeps

While differences in gene content may result in the evolution of overlapping ecotypes or eventually to sympatric speciation, the higher rates of recombination in some of the North Pond populations should produce evidence of gene-specific sweeps. Likely due to the increased cutoff values used to determine the location of SNVs, compared to Crits-Christoph *et al*. (2020), we did not observe a correlation between nucleotide diversity and gene coverage or purifying selection. As such, we used nucleotide diversity from the six MAG populations present over multiple time points as a proxy for potential gene sweep events based on a statistically significant decrease in nucleotide diversity for individual genes compared to the genomic mean in all time points (Table 3; Supplemental Data 7-9).

**Table 3.** Number of genes with decreased nucleotide diversity and regions detected with variable F_ST_.

| Genome | No. of CDS | No. of low nucleotide diversity CDS° | Low dN/dS (< mean genome value)* | dN/dS > 1* | no dN/dS* | Longest consecutive low nucleotide diversity CDS | No. of elevated $F_{ST}$ regions (> 1 std dev) | No. of elevated $F_{ST}$ regions (> 2 std dev) |
| --- | --- | --- | --- | --- | --- | --- | --- | --- |
| NORP83 | 2,657 | 60 (2%) | 4 (7%) | 9 (15%) | 43 (72%) | 4 | 8 | 0 |
| NORP139 | 1,538 | 32 (2%) | 6 (19%) | 12 (38%) | 10 (31%) | 1 | 0 | 0 |
| NORP147 | 2,650 | 0 (0%) | 0 (0%) | 0 (0%) | 0 (0%) | 0 | 0 | 0 |
| NORP163 | 2,182 | 190 (9%) | 37 (19%) | 11 (6%) | 117 (62%) | 5 | 14 | 1 |
| NORP169 | 2,842 | 238 (8%) | 19 (8%) | 19 (8%) | 175 (74%) | 3 | 3 | 0 |
| NORP246 | 3,851 | 22 (1%) | 5 (23%) | 3 (14%) | 12 (55%) | 2 | 0 | 0 |

°Percentage calculated as no. of low nucleotide diversity CDS/total CDS. \*Percentage calculated as fraction of CDS with low nucleotide diversity

Four of the MAG populations had a small percentage of putative coding sequences (0-2%) that could represent gene-sweep events (Table 3). Both NORP83 (2%) and NORP147 (0%) were previously identified as having low predicted rates of recombination. The low number of putative gene sweeps is likely a direct result of low recombination rates, as low rates of recombination would increase gene linkage and thus prevent individual genes from sweeping through the population. However, both NORP139 (2%) and NORP246 (1%) had small percentages of their respective coding sequences that may have represented gene-specific sweeps, but estimates of recombination rates for these populations was higher compared to NORP83, NORP147, and NORP169. Conversely, NORP163 (9%) and NORP169 (8%) had a larger percentage of genes involved in putative gene-sweep events, though NORP163 was identified as having a relatively low recombination rate. The contradiction between a low expected recombination rate and a larger number of putative gene-sweeps is likely the result of how the four-gamete test and inStrain recombination rates are calculated. Both approaches use linked biallelic SNVs to estimate recombination rates, but recent gene-sweeps, especially sweeps through multiple strains, would not have many SNVs. With the exception of NORP139, most of the putative gene-sweeps (55-74%) did not produce a dN/dS value from the Gretel approach analyzing all potential haplotypes and isoforms (Table 3; Supplemental Data 9). A smaller percentage of putative gene-sweeps were attributed to coding regions under purifying selection (7-23%; gene_dN/dS_ *<* mean genome_dN/dS_) and relaxed selection (6-38%; dN/dS *>* 1) (Table 3; Supplemental Data 9). While this might suggest that most genes are not under selection, this methodology filtered out putative coding regions without multiple haplotypes/isoforms, lacking mutations (*dN* = *dS* = 0), and with short timescales since divergence (dS *<* 0.01). These data could support the interpretation that the identified putative gene-sweeps are recent and have yet to accumulate mutations or been acted upon by selection. Given the inferred slow growth rates among the North Pond microbial community, longer periods of time may be required to observe such mutations and selection in a population.

We also searched for evidence of genetic differentiation among subpopulations using the pairwise fixation index F_ST_ by identifying genomic regions of 5+ genes that had variable allele frequencies across populations (Supplemental Data 10). Generally, this metric followed the same pattern observed using nucleotide diversity. NORP163 and NORP169 had several regions with elevated F_ST_ when considering a deviation greater than one standard deviation from the mean F_ST_ value. Despite its low estimated recombination rate, NORP83 had the second highest number of regions with increased F_ST_ (*n* = 8). For all three genomes, none of the elevated F_ST_ regions shared gene content with the putative gene-sweep regions identified with nucleotide diversity. However, increasing the required deviation to greater than two standard deviations from the mean reduced the number of differentiated regions to a single region in the NORP163 population. Due to the constraints placed on calculating the F_ST_ metric, which requires genes in two time points to have corresponding biallelic SNVs, many regions were likely excluded from this F_ST_ analysis. Additionally, the F_ST_ calculations required at least five genes in a variable region, though results from the nucleotide diversity metric demonstrated that the longest stretch in all genomes of decreased diversity was five sequential genes (Table 3). Further, the nucleotide diversity metric could be calculated for all genes and the corresponding reads for all time points that met the inStrain cutoffs. Very few genomic regions had significantly elevated F_ST_ results, especially when filtered using a more stringent cutoff, supporting that even though some putative gene-specific sweeps were detected in the populations, the observed low recombination rates likely limit the total number of events between strains.

Collectively, for all six of these populations across all time points, there were several KEGG functional BRITE categories that were over represented within the identified putative gene-sweep regions compared to the genomic averages, including tRNA biogenesis (BR03016), Ribosome (BR03011), Ribosome biogenesis (BR03009), Transcription factors (BR03000), and Chaperones and folding catalysts (BR03110) (Supplemental Data 11). On average, all of these categories, except Transcription factors, also had increased mean dN/dS values in the putative gene-sweep regions (0.67-1.69) compared to the genomic average (0.40-0.48). Though the number of putative gene-sweep regions only reflects a small percentage of the total genes in the six MAGs, it is interesting that several core functional categories have comparatively relaxed selection. This could imply that the accumulation of mutations within core functional genes does not impart a fitness cost, which could again be a result of the slow growth conditions experienced by these populations. From the perspective of all six populations, the system dynamics appear to have been molded by dispersal of strains across the aquifer, and the low to moderate rates of homologous recombination may have facilitated species cohesion and gene transfer among closely related populations.

## Conclusion

The MAGs described here represent subseafloor microbial populations that exhibited substantial changes over the ∼825-day sampling period. Thus, these datasets provide evidence that the marine crustal aquifer hosts a dynamic habitat in which microbial populations grow and decay and can be dispersed over monthly timescales. While rapid allele frequency shifts can be linked to different types of population interactions, such as sweeps, dispersal, and clonal expansion, dispersal appears to play an important role in structuring the most abundant populations in the crustal fluid samples. The shifts in allele frequencies and nucleotide diversity suggest that stochastic events, such as dispersal and the mixing of populations throughout the aquifer, also mold the evolutionary trajectories of microbial populations in this habitat and that selection and drift occur on timescales that were not captured as part of this study (36). Though classically structured as the introduction or removal of mutations through selection and drift, evolution, broadly defined, reflects changes in allele frequency within a population. Thus, dispersal can introduce extant populations from elsewhere in the environment and the interaction of these newly introduced alleles with the environment reflects an evolutionary response. As such, we are able to observe microbial evolution in action in the fluids moving through the oceanic crust. The observed population shifts occur in the span of months, reflecting the highly dynamic nature of the system, both biologically and physically. The accumulation of extra-chromosomal insertions, such as transposable elements and plasmids, suggests that relaxed selection pressures and slow generation times can impact these populations in a manner distinct from dispersal. In summary, the subseafloor aquifer of North Pond represents a highly dynamic habitat where evolution may be governed largely by the stochastic forces of dispersal.

## Materials and Methods

### Sample collection and sequencing

As has been described elsewhere (6, 19, 22), North Pond water samples were collected from two CORKs at holes U1382A and U1383C in 2012 and 2014. For sample time points (TP) during expeditions (TP0 in April 2012 and TP9 in April 2014), the Mobile Pumping System (MPS) attached to the *ROV* Jason II was used to collect samples through umbilical lines connecting the CORK platform to fluids in the aquifer. For samples collected between expeditions (TP1-TP8), the battery-powered GEOMicrobeSled (37, 38) deployed on the CORK collected samples approximately every two months. In all instances, lines were purged to remove stagnant water prior to sampling and all samples were fixed with RNALater in situ. A complete set (*n* = 8) of *in situ* samples were collected from U1382A and provide the necessary samples and resolution for downstream analysis (as described below). Previously described in Tully *et al*. (2018), DNA was extracted from filters using a phenol chloroform method (39) and paired-end sequencing was performed on an Illumina HiSeq 1000 at the Marine Biological Laboratory. Raw sequences were quality controlled using Cutadapt (40) v1.7.1 (parameters: `-e 0.08 --discard-trimmed --overlap = 3`) and Trimmomatic (41) v0.33 (parameters: `PE SLIDINGWINDOW:10:28 MINLEN:75`).

### Metagenome-assembled genomes and manual curation

To build upon the set of MAGs generated in Tully *et al*. (2018), the quality controlled paired-end reads were assembled using three different approaches to maximize recovery of MAGs that had not previously been assembled into MAGs. A detailed methodology has been provided in the SI Appendix and a comprehensive assessment of the final set of MAGs can be found in Supplemental Data 1.

### SNV identification and analysis

anvi’o (42, 43) v5.0 contig databases were generated for all MAGs. FASTA files of the contigs of each MAG were converted to an anvi’o database (`anvi-gen-contigs-database --skip-mindful-splitting`). Bowtie2 (44) v2.3.4.1 (parameters: `--no-unal`) was used to create recruitment profiles for each MAG in each sample. The output SAM format file was converted to a sorted BAM format file using samtools (45) v1.9 and reads with *<* 95% sequence identity over 75% of the alignment were filtered from the recruitment profile using BamM v1.7.3 (parameters: `--percentage_id 0.95 --percentage_aln 0.75`; https://github.com/Ecogenomics/BamM). Recruitment profiles were merged with `anvi-merge`. During the `anvi-profile` step, the flag `--profile-SCVs` was set to include identification of single nucleotide variants (SNVs) and single codon variants (SCVs). Each individual MAG anvi’o database had the collection value ‘DEFAULT’ and bin value ‘EVERYTHING’ added using `anvi-script-add-default-collection`. Using those collection and bin values, `anvi-gen-variability-profile` (default parameters; `--engine AA --include-contig-names --engine CDN`) to extract SNV, single amino acid variants (SAAVs), and SCVs, from the profiles of each MAG in each North Pond sample (46) (Supplemental Data 3). The SCV and SAAV values were used to calculate pN/pS ratios using `anvi-script-calculate-pn-ps-ratio` (anvi’o v6.2). The sorted, filtered BAM format recruitment profiles described above were used to determine read counts for each contig using featureCounts (47) v1.5.3 (default parameters) as implemented within Binsanity-profile (48) v0.3.3 (default parameters). Read counts were converted to the normalized unit reads per kilobase pair MAG per Megabase pair of metagenomic sample (RPKM; Supplemental Data 2). Read counts were also used to determine the relative fraction of each MAG in each metagenomic sample.

Hole U1382A provided a high-resolution sample dataset (10 time points over ∼825 days), and MAGs detected in these samples were selected for downstream analysis based on two criteria from the time series: (1) sufficient coverage (≥5 RPKM) in ≥3 time points or (2) high coverage (≥30 RPKM) in at least one time point. Only time points that had ≥5 RPKM were considered for SNV calculations. This criterion was set to ensure that most base pairs in each MAG had at least 20× read coverage, which was the minimum coverage value to determine SNVs in all downstream analyses (see below).

As has been reported previously (26, 49, 50), MAGs tend to represent a cohesive population represented by a >95% nucleotide identity boundary. To confirm that a majority of the signal for each population came from closely related organisms, RPKM values were recalculated and compared using recruited reads filtered to 99% identity over 75% of the length of the alignment using BamM (parameters: `--percentage_id 0.99 --percentage_aln 0.75`; Supplemental Data 2). Additionally, results from inStrain (51) were used to track what fraction of reads recruited at 95-100% identity (see below).

Two of the MAGs (NORP143 and NORP246) were taxonomically assigned to the genus *Shewanella* and shared 98.2% ANI, as determined with FastANI (49). For consideration of this analysis these MAGs were deemed to represent the same species-level population and NORP143 (67.24% complete, 9.87% redundancy) was removed from downstream analysis while NORP246 was retained (82.57% complete, 2.28% redundancy; Table 1). The features identified in NORP246 were confirmed by creating a new anvi’o database with recruited reads filtered to 99% identity over 75% of the length of the alignment (as above). To ensure consistency, analyses using anvi’o calculated values below were based on recruited reads filtered to 95% identity over at least 75% of the length of the alignment.

### Functional annotation and taxonomic assignment

Protein sequences as determined within anvi’o were submitted to the GhostKOALA KEGG annotation and mapping service (52) (Submitted 23 October 2018; parameters: `genus_prokaryotes` + `family_eukaryotes`). KEGG annotations were added to the anvi’o contig databases for each MAG using the Python script `KEGG-to-anvio` (https://github.com/edgraham/GhostKoalaParser) and `anvi-import-functions` (http://merenlab.org/2018/01/17/importing-ghostkoala-annotations/).

GTDB-Tk (53) v1.3.0 (`classify_wf` default parameters) using database R95 was used to determine a taxonomic assignment for each MAG.

### Data transformations for analysis

The SNV occurrence table generated by anvi’o was filtered by retaining all entries with *>* 0 entropy value, SNV positions ≥ 20× coverage in the time point(s) of interest, and if the departure from consensus was ≥10%. Putative SNVs at each time point were converted to single nucleotide variants per kilobase pair (SNVs kbp^-1^) (23), a measure of polymorphisms within the population, based on the full length of the MAG (Supplemental Data 3). Time points for which a MAG was present at *<* 5 RPKM were not used to calculate SNVs kbp^-1^ or considered in further analysis.

We used anvi’o to obtain metrics of microbial population dynamics based on MAGs. anvi’o is a widely used platform for microbial genomics and provides a baseline to determine polymorphisms in the underlying metagenomic reads. The criteria detailed here provide a conservative method to compare features such a SNV frequency, major allele frequency, and gene coverage. For data processed with anvi’o, major allele frequency was determined by counting the frequency of nucleotides in the aligned reads, identifying the consensus allele, and dividing the number of occurrences of that allele by the total coverage for that site (*i*.*e*., the nucleotide in the reference MAG sequence was not considered in this calculation). For each MAG, mean major allele frequency was determined by averaging all major allele frequencies for each SNV across the entire MAG. For comparing time points, if a SNV was detected in one time point, but not another (and coverage ≥20×), the positions for which no SNV was recorded were assumed to be fixed at the consensus allele. Hierarchical clustering was performed and plotted using seaborn (doi: 10.5281/zenodo.3767070) with the default clustermap settings (average distance linkage method and Euclidean distance metric).

### Strain Identification and Abundance

DESMAN (54) was used to identify strains of MAGs based on SNV proportions in core genes. DESMAN offers insight into the estimated complexity of the populations captured at a time point for a particular MAG. This analysis calculates the ratio of alleles at each time point in a set of core genes in order to identify discrete haplotypes (referred to throughout as “strains”) and to define an estimate of relative strain contribution to the total population. This disentangles the bulk signal provided by the allele frequencies and offers a hypothesis as to how and why specific alleles are changing. We used anvi’o to first identify clusters of orthologous genes (COGs) in the MAGs, as well as to identify SNV variants in a set of 36 universal single-copy COGs as defined by (55). The DESMAN haplotype identifier was computed five times with a varying number of haplotypes (2-8), and the posterior mean deviance was plotted to select the optimal haplotype number and best replicate run. Results from Gretel (56) were used to support the optimal haplotype number for each MAG when running DESMAN (see below). Only time points with sufficient coverage (as described above) were included in depictions of haplotype abundance across samples.

### Recombination

We used both mcorr (27) and inStrain (51) to compile further metrics of population dynamics and to assess recombination rates. The software package mcorr was used to determine the recombination rate for the populations represented by the MAGs. mcorr utilizes a correlation matrix to link recombination events at the 300 bp scale and fits these events to a model in order to estimate recombination and mutation rates. The correlation matrix requires polymorphisms in close proximity to correlate in an expected manner, which can be confounded as the complexity of the underlying subpopulation increases and rate recombination increases between subpopulations. Additionally, should the population in question not adhere to the model trained from laboratory experiments, estimates of recombination and mutation may not be accurate or recoverable. An mcorr correlation profile for each time point was constructed using filtered recruitment profiles generated with the script `filter_reads.py` (parameters: `-m 0.96 -q 2 -l 1500`; https://github.com/alexcritschristoph/soil_popgen/blob/de1e09dd8416131a6b5feba74f0c3a5747d38da1/inStrain_lite/inStrain_lite/filter_reads.py) and a GFF formatted gene prediction file generated by Prodigal (57) v2.6.3 (parameters: `-p m -m`) using `mcorr-bam` (default parameters). The correlation profiles were fit to the mcorr model using `mcorr-fit` (default parameters). The correlation profiles and calculated fit were assessed for clear evidence of monotonic decay and normalized distribution of residuals. The estimated ratio of recombination-to-mutation (gamma/mu; *γ/µ*) was incorporated into further analysis if the bootstrapping mean was *<* 2× the estimated value (24) (Supplemental Data 4). Values of gamma/mu that failed this cutoff were interpreted as evidence of recombination amongst multiple haplotypes that could not be resolved using the correlation profile approach.

In contrast to mcorr, the inStrain profile links polymorphisms over the length of the metagenomic read. Thus, complex populations with extensive recombination, which may have been missed through the use of the mcorr correlation matrix, could be resolved but limited by the length of the read. Along with the accompanying genome statistics, inStrain provides highly precise measures of the metapopulation. inStrain was used to profile the underlying diversity on a genome-wide basis for each MAG and on a gene-by-gene basis using Prodigal predicted ORFs from each time point (inStrain profile; parameters: `-l 0.95 --min_mapq 1 -c 20 -f 0.1`). Parameter settings, including read percent identity (95%), coverage (20×), and minimum allele frequency (10%), were selected to ensure that SNV detection by inStrain and anvi’o were conducted with the same level of stringency. inStrain includes the ability to filter reads that mapped to multiple locations during recruitment and these reads were removed, if present. Output from inStrain was assessed for the presence of linkage disequilibrium (*r*^*2*^), the distribution of read recruitment identity, per gene and genome average nucleotide diversity (*π*), which provides a probability for two reads having the same nucleotide at a particular position, and estimate of replication rate (51) (Supplemental Data 4). Linkage decay, a signal of recombination, was observed by plotting *r*^*2*^ against the distance between linked SNVs; where distance between linked SNVs were grouped into 10 bp ranges and separated by the type of mutation within the coding region (synonymous-synonymous [S-S], synonymous-nonsynonymous [S-N], and nonsynonymous-nonsynonymous [N-N]; Supplemental Data 5). Additionally, the four-gamete test, another signal of recombination, was performed for all biallelic (AB and ab haplotypes) linked SNV locations (11). Under the infinite-site model, the presence of all four haplotypes (AB, ab, Ab, aB) can only be explained by at least one recombination event (58, 59). The frequency of the occurrence of the number of haplotypes (one, two, three, or four haplotypes present) between two sites was calculated. Additionally, to identify putative genes that had undergone a selective sweep by detecting genes with statistically significantly lower nucleotide diversity, the mean MAG nucleotide diversity was compared to the gene mean for all time point of interest with the Welch’s t-test (Supplemental Data 7). A corresponding dN/dS value was calculated, when applicable, as computed by Gretel (see below).

The SNV and linkage output from inStrain was used to compute F_ST_, a measure of allele frequency differences between two populations. For all linked biallelic sites within putative coding regions that had a complete start and stop codon, a moving Hudson F_ST_ calculation (60, 61) was performed as implemented in the scikit-allel package v1.3.3 (https://scikit-allel.readthedocs.io/) consistent with Crits-Christoph *et al*. (2020). Genes with coverage outside of two standard deviations of the genome mean were excluded. A pairwise calculation was performed between all time points of interest and a genome mean was reported. All remaining putative coding sequences were screened in a five gene window for elevated an F_ST_ signal that exceeds either one or two standard deviations from the genome mean (Supplemental Data 10).

### Gene frequency

For each MAG, gene coverage values were exported from the anvi’o databases using `anvi-export-gene-coverage-and-detection`. Gene frequency in a sample was calculated by dividing each gene coverage value by the median coverage value of all genes. Genes of interest were identified based on changes in gene frequency such that the maximum frequency value minus the minimum frequency value in the time points of interest was ≥1. Genes identified in this manner lacking a KEGG annotation were compared against GenBank nr (62) using BLASTP (63) (Supplemental Data 6). Hierarchical clustering was performed and plotted using seaborn (as above).

### Gene variants

The software package Gretel (56) was used to reconstruct haplotypes on a gene-by-gene basis (in contrast to DESMAN, which recovers haplotypes or strains for whole genomes based on SNVs in core genes). Gretel greedily uses the polymorphism patterns of the recruited reads to reconstruct all possible haplotypes for the genes within a MAG. This approach can lead to chimeric reconstructed genes, as clearly demonstrated by the presence of haplotypes within internal stop codons, but also offers the full potential space of haplotypes for a gene within a time point. We simplified that potentially inflated value by converting the haplotypes to isoforms; this constrained estimate and describes a reduced space of environmental proteins. The combined mean dN/dS for each gene extended our ability to assess selection within the population. The filtered recruitment profiles for each MAG and time of interest was used to construct a variant call format (VCF) file using `gretel-snpper` (default parameters), compressed with `bgzip` (default parameters), and indexed with `tabix` (default parameters). Gene positions on each contig were determined from the Prodigal gene predictions and used as inputs for the analysis by `gretel` (default parameters). If a gene was determined to be in the reverse direction, putative haplotypes were reverse complemented and all correctly oriented genes were translated to proteins using seqmagick v0.8.4 (https://fhcrc.github.io/seqmagick/). Haplotypes that translated to isoforms with internal stop codons were excluded from further analysis. Haplotypes and corresponding isoforms for each putative gene from all time points of interest were combined (Supplemental Data 8). CD-HIT (64) was used to cluster haplotypes and isoforms with 100% identity (parameters: `-c 1 -d 0 -g 1`). The combined isoforms were aligned using MUSCLE v3.8.31 (65) (default parameters) and the alignments were used as inputs to PAL2NAL (66) (default parameters) to create a codon alignment of the corresponding haplotypes. dN, dS, and dN/dS was calculated (67) for all genes with detected haplotypes using PAML (68) (example control file: Supplemental Information; Supplemental Data 9). The output was filtered to include only dN > 0, 0.01 < dS < 1, and dN/dS < 5. dN/dS and dS for all pairwise haplotype comparisons were averaged. The distribution of the number of isoforms predicted for each gene was used to inform the selection of target strains in the DESMAN analysis.

## Data Availability

Data for this project can be found deposited at DDBJ/ENA/GenBank under the BioProject accession no. PRJNA391950. Raw sequence reads are available through the Short Read Archive with the accession no. SRX3143886-SRX3143902. Raw sequence reads from Meyer *et al*. (2016) constituting the metagenomic samples from 2012, are available under the BioProject accession no. PRJNA280201. Additional files are available through figshare, including the contigs, anvi’o protein calls, Prodigal protein calls, anvi’o database profiles for the MAGs (doi: 10.6084/m9.figshare.17698631), and the filtered BAM files used in this analysis (doi: 10.6084/m9.figshare.17701254). As detailed in Supplemental Data 1, all new/updated MAGs have been submitted to NCBI under the BioProject accession no. PRJNA391950. Scripts created to process data, perform analyses, and create plots have been made accessible through GitHub (https://github.com/bjtully/northpond_evolution).

## ACKNOWLEDGMENTS

We thank the crews of the *R/V* Merian and *ROV* Jason II, Wolfgang Bach, Peter Girguis, Chih-Chiang Hsieh, Ulrike Jaekel, Beate Kraft, Huei-Ting Lin, Beth Orcutt, Keir Becker, Stephanie Carr, and Heiner Villinger for support in accomplishing the 2012 and 2014 field programs. Ship time was provided by the German Science Foundation (DFG). Both Katrina Edwards and James Cowen’s efforts were critical to the field component and success of this project. They are missed immensely. Leslie Murphy and Emily Reddington provided laboratory support at the WM Keck sequencing facility at the Marine Biological Laboratory. This work was supported by NSF OCE-1062006, OCE-1745589 and OCE-1635208 to J.A.H. The Gordon and Betty Moore Foundation sponsored observatory components at North Pond through grant GBMF1609. The Center for Dark Energy Biosphere Investigations (C-DEBI) (OCE-0939564) supported J.A.H. and B.J.T. This is C-DEBI contribution XXX.

