## Supplemental Information for "Microbial populations are shaped by dispersal and recombination in a low biomass subseafloor habitat"

#### This PDF file includes:

Supplementary text
Figs. S1 to S2
Tables S1 to S2
Legends for Dataset S1 to S11
SI References

#### Other supplementary materials for this manuscript include the following:

Datasets S1 to S11

### Supporting Information Text

### Methodology

To build upon the set of MAGs generated in Tully *et al.* (2018)(1), the quality controlled paired-end reads were assembled using three different approaches to maximize recovery of MAGs that had previously remained unbinned. Method 1 applied an approach developed as part of Tully *et al.* (2018). Each individual sample (n = 21) from the 2012-2014 North Pond metagenome dataset was assembled using Megahit (2) (v1.1.2; parameters: `-presets meta-sensitive`). CD-HIT-EST (3) (v4.6; parameters: `-T 90 -M 5000000 -c 0.99 -n 10`) was used to cluster contigs with ≥99% nucleotide identity for all contigs ≥2kb from all samples. This set of primary contigs were then co-assembled using Minimus2 (AMOS v3.1.0; parameters: `-D OVERLAP=100 MINID=95`) (4) to produce a set of secondary contigs. Secondary and remaining primary contigs ≥3kb were binned using BinSanity (5) using a custom Binsanity-lc workflow (v0.2.6; parameters: Table S1; https://github.com/edgraham/NorthPondBinning_ScriptRepo). Briefly, BinSanity was run iteratively for six passes, where after each pass high completion bins (≥50% complete) were identified using CheckM (v1.1.1; parameters: `lineage_wf` default parameters) (6) and contigs contributing to the bins were removed from additional passes. For pass seven, all bins that had been identified as "high redundancy" (≥50% complete, ≥10% redundancy) were refined using Binsanity-wf (v0.2.6). Bins determined to be ≥50% complete and <10% redundancy were designated as MAGs for downstream analysis. Preference and k-means values used for each pass are available in Table S1.

Method 2 applied a subsampling approach, whereby reads from each sample were randomly subsampled to 1-10, 12, 15, 17, 20, 30, and 50% of the total paired end reads using seqtk (v1.2; parameters: `sample, -s random seed between 1-300`; https://github.com/lh3/seqtk). Thus, each sample was split into 16 different paired end read datasets. Each subsampled dataset was assembled using Megahit, as above. The primary contigs from the subsampled dataset were combined with the contigs generated from the full read dataset using CD-HIT-EST and Minimus2, as above. Remaining primary and secondary contigs ≥3kb were binned as above. Method 3 was a co-assembly combining all 21 paired end datasets using Megahit, as above. Contigs ≥3kb were binned, as above. Additionally, the original set of contigs generated in Tully *et al.* were re-binned using the Binsanity-wf (v0.2.6; default parameters).

New MAGs were compared to the original 195 NORP MAGs using FastANI (7) (v1.0; parameters: `-fragLen 1500`). Clusters of MAGs with ≥98.5% average nucleotide identity were considered identical and a representative with highest quality score, determined as completion - (5 $\times$ redundancy), was selected. In this manner, the new MAGs could revise an original MAG if the overall quality statistics improved. The highest quality MAG was submitted to NCBI (see Data Availability).

### Results and Discussion

**Improving North Pond metagenome-assembled genomes.** Many times, metagenomic datasets are treated as static elements that are mined only once. But as understanding about the underlying algorithms and dataset complexity continue to advance, there is an opportunity to extract additional information from previously analyzed metagenomes. This reanalysis of the North Pond 2012-2014 metagenomic time series capitalizes on the fully released BinSanity binning software tool and increased clarity regarding how microdiversity within an metagenomic sample can impact contig recovery at the assembly step, preventing the generation of high-quality MAGs. In Tully *et al.* (2017) (1), metagenomic samples were co-assembled based on sample origin, generating 78,004 contigs ≥3kb (N50 25,932 bp) and the binning method generated 195 high-completion MAGs (Table S2). The amended three methods applied to the dataset generated more contigs ≥3kb (132,025-170,001 contigs) with lower N50 (11,653-19,010 bp), but on average generated more high-completion MAGs (140-231 MAGs; Supplemental Data 1).

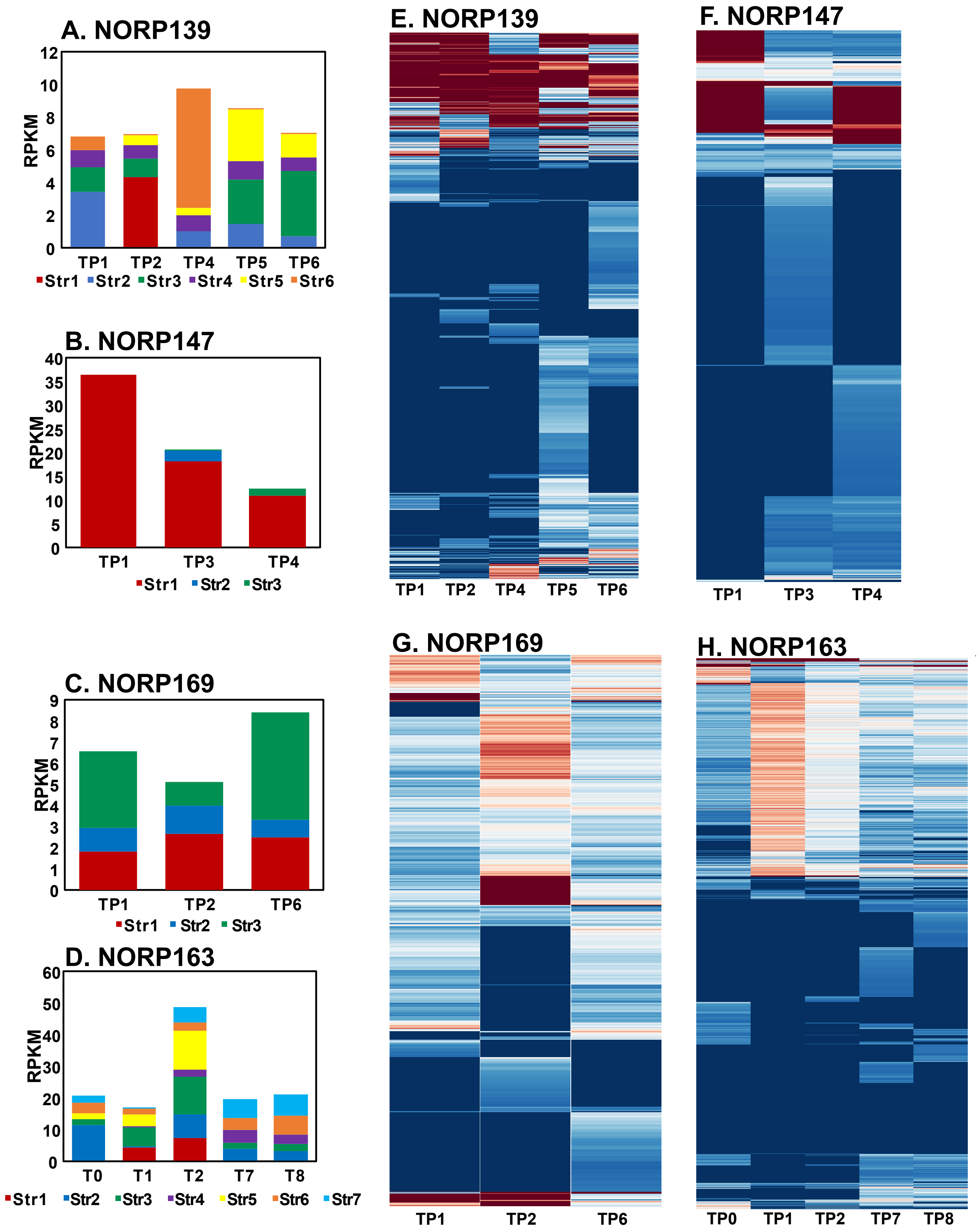


**Fig. S1.** Strain relative abundance as determined using DESMAN for (A) NORP139, (B) NORP147, (C) NORP169, and (D) NORP163. Major allele frequency for SNVs detected in time points of interest for (E) NORP139, (F) NORP147, (G) NORP169, and (H) NORP163. The major allele frequencies have been hierarchically clustered and scaled from 0-1. Both 0.0 and 1.0 represent a fixed allele; 0.5 represents a split allele.

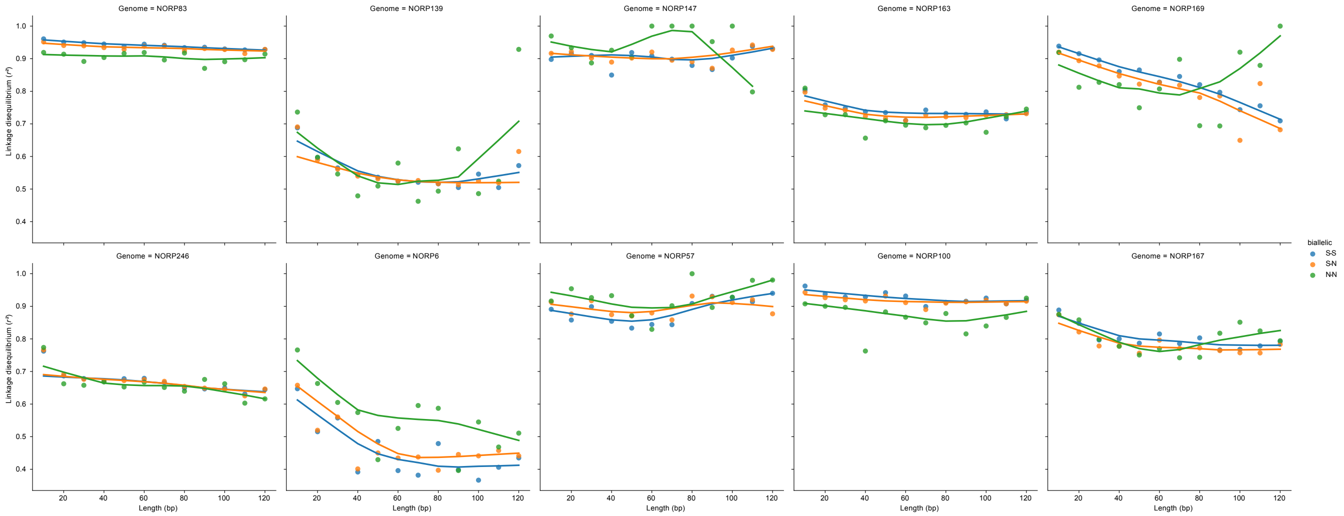


**Fig. S2.** Linkage disequilibrium of $r^2$ for linked SNVs pairs for all MAGs. Each circle is the mean $r^2$ for pairs of linked SNVs at that distance range (*e.g.,* 1-10 bp, 11-20 bp, *etc.*). Linked SNVs are denoted by their predicted mutation type (nonsynonymous: N, synonymous: S).

**Table S1. Settings used for each of the 7 passes with Binsanity-lc.**

| Pass | Preference | Refinement preference | K-means value |
|---|---|---|---|
| 1 | -25 | -50 | 500 |
| 2 | -15 | -35 | 50 |
| 3 | -10 | -25 | 50 |
| 4 | -5 | -25 | 50 |
| 5 | -3 | -25 | 50 |
| 6 | -2 | -25 | 50 |
| 7 | -2 | -10 | NA |

**Table S2. Assembly statistics comparing original 2017 result with modified assembly and binning protocols presented here.**

| Method | BinSanity approach | No. of contigs $\geq$3kb | No. of contigs $\geq$100kb | N50 | No. of MAGs $\geq$50 | No. of MAGs $\geq$90 |
|---|---|---|---|---|---|---|
| Tully et al. (2018) | pre-release BinSanity | 78,004 | 1,278 | 25,932 | 195 | 68 |
| Tully et al. (2018) | Binsanity-wf | 78,004 | 1,278 | 25,932 | 222 | 67 |
| Method 1 | Binsanity-lc | 103,964 | 1,048 | 19,010 | 209 | 96 |
| Method 2 | Binsanity-lc | 170,001 | 1,498 | 17,806 | 140 | 46 |
| Method 3 | Binsanity-lc | 132,025 | 612 | 11,653 | 231 | 45 |

**SI Dataset S1** (**SupplementalData1-NPMAGStats.xlsx**)

MAG quality values with details on relationship to original Tully *et al.* (2018) MAGs.

**SI Dataset S2** (**SupplementalData2-RPKM.xlsx**)

MAG RPKM values in all time points of interest using variable identity cutoffs in the BamM step. Each MAG is also assessed for the number of samples it is "present" in using different RPKM cutoff values.

**SI Dataset S3** (**SupplementalData3-SNVs-NucDiv-iRep.xlsx**)

MAG values for total SNVs, length, SNV per kbp, nucleotide diversity, and T:P ratio.

**SI Dataset S4** (**SupplementalData4-Summarized-inStrain-Genome-Results.xlsx**)

inStrain results for each genome by time point of interest.

**SI Dataset S5** (**SupplementalData5.tar.gz**)

The set of inStrain `linkage.csv` and `SNVs.tsv` used to calculated linkage disequilibrium.

**SI Dataset S6** (**SupplementalData6-GeneFrequencyValuesAnnotations.xlsx**)

For each MAG, the genes that were determined to have variable coverage across time points. Gene IDs and coverage values determined from anvi'o output. Top two BLAST matches in the NR database are provided for each gene and KEGG assignment (when available).

**SI Dataset S7** (**SupplementalData7-Putative-Gene-Sweeps.xlsx**)

For each MAG, genes determined to be have statistically significant differences in nucleotide diversity, as a proxy for putative genes that have undergone a selective sweep. Significance determined by Welch's t-test. Mean dN/dS was determined from Gretel results.

**SI Dataset S8** (**SupplementalData8-GretelResults.tar.gz**)

Output for the Gretel analysis. There is a directory for each gene from each MAG with the raw Gretel results.

**SI Dataset S9** (**SupplementalData9-PAMLResults.tar.gz**)

Output for the PAML analysis There is a directory for each gene from each MAG based on the Gretel results.

**SI Dataset S10** (**SupplementalData10-FST.xlsx**)

Reports the population mean FST values for genes that met cutoff criteria. Calculated in a pairwise manner between time points of interest. Reports the genes and region number when 5+ genes were detected with mean FST >1 standard deviation above the genome mean. Reports the genes and region number when 5+ genes were detected with mean FST >2 standard deviation above the genome mean. When relevant includes corresponding data from Supplemental Data 7. Performed in a pairwise manner between time points of interest. Reports num. of SNV per gene, FST, p, and coverage.

**SI Dataset S11** (**SupplementalData11-KEGGCategories.xlsx**)

Provides information on the KEGG categories of genes determined to have putative selective sweeps.
